# Gene model for the ortholog of *DENR* in *Drosophila eugracilis*

**DOI:** 10.64898/2026.06.23.734050

**Authors:** Megan E. Lawson, Kylee Sanow, Ines Martinand, Mihai Fratian, Madelyn Matura, Chinmay P. Rele, Laura K. Reed, Jeffrey S. Thompson, Kellie S. O’Rourke

## Abstract

Gene model for the ortholog of *Density regulated protein* (*DENR*) in the Apr. 2013 (BCM-HGSC/Deug_2.0) (DeugGB2) Genome Assembly (GenBank Accession: GCA_000236325.2) of *D. eugracilis*. This ortholog was characterized as part of a developing dataset to study the evolution of the Insulin/insulin-like growth factor signaling pathway (IIS) across the genus *Drosophila* using the Genomics Education Partnership gene annotation protocol for Course-based Undergraduate Research Experiences.

## Introduction

“Computational gene predictions in non-model organisms often can be improved by careful manual annotation and curation, allowing for more accurate analyses of gene and genome evolution (Mudge and Harrow 2016; Tello-Ruiz et al., 2019). The Genomics Education Partnership (thegep.org) uses web-based tools to allow undergraduates to participate in course-based research by generating manual annotations of genes in non-model species (Rele et al., 2023). These models of orthologous genes across species, such as the one presented here, then provide a reliable basis for further evolutionary genomic analyses when made available to the scientific community. The particular gene ortholog described here *Density regulated protein* (*DENR*) in *D. eugracilis* was characterized as part of a developing dataset to study the evolution of the Insulin/insulin-like growth factor signaling pathway (IIS) across the genus Drosophila.” (Myers et al., 2024).

“The IIS pathway is a highly conserved signaling pathway in animals and is central to mediating organismal responses to nutrients (Hietakangas and Cohen 2009; Grewal 2009)” (Myers et al., 2024). “*DENR* was first discovered in a human teratocarcinoma cell line because its concentration in cells increased with cell density (Deyo et al., 1998). Subsequent bioinformatic and biochemical analyses showed that the protein is conserved across eukaryotes and functions in non-canonical translation initiation (Fleischer et al., 2006; Skabkin et al., 2010). *D. melanogaster* flies homozygous for a null, knockout allele of the gene encoding *DENR* (FBgn0030802), die as pharate adults, showing a larval-like epidermis and reduced proliferation of histoblast cells (Schleich et al., 2014). Subsequent experiments using both RNAi in S2 cells and the knockout allele in larvae showed that DENR is required, along with its interacting partner MCT-1, for the proper expression regulation of a subset of transcripts required for cell cycle progression and growth. In particular, the loss of *DENR* reduces expression of the insulin receptor and makes larvae less sensitive to insulin signaling (Schleich et al., 2014), thus implicating DENR in the regulation of the insulin signaling pathway.” (Laskowski et al., 2024).

“*D. eugracilis* is part of the melanogaster species group within the subgenus *Sophophora* of the genus *Drosophila* (Pélandakis et al., 1993). It was first described as *Tanygastrella gracilis* by Duda (1924) and revised to *Drosophila eugracilis* by Bock and Wheeler (1972).

*D. eugracilis* is found in humid tropical and subtropical forests across southeast Asia (https://www.taxodros.uzh.ch, accessed 1 Feb 2023).” (Morgan et al., 2022).

We propose a gene model for the *D. eugracilis* ortholog of the *D. melanogaster Density regulated protein* (*DENR*) gene. The genomic region of the ortholog corresponds to the uncharacterized protein XP_017064488.1 (Locus ID LOC108103491) in the Apr. 2013 (BCM-HGSC/Deug_2.0) (DeugGB2) Genome Assembly of *D. eugracilis* (GenBank Accession: GCA_000236325.2). This model is based on RNA-Seq data from *D. eugracilis* (Chen et al., 2014; PRJNA63469) and *DENR* in *D. melanogaster* using FlyBase release FB2024_02 (GCA_000001215.4; Gramates et al., 2022; Jenkins et al., 2022; Larkin et al., 2021). The Genomics Education Partnership maintains a mirror of the UCSC Genome Browser (Kent WJ et al., 2002; Gonzalez et al., 2021), which is available at https://gander.wustl.edu.

## Results

### Synteny

The target gene, *DENR*, occurs on chromosome X in *D. melanogaster* and is flanked upstream by *CG4880* and *CG13002* and downstream by RNA polymerase III subunit I *(Polr3I)* and Nitrogen permease regulator-like 2 *(Nprl2)*. The *tblastn* search of *D. melanogaster* DENR-PA (query) against the *D. eugracilis* (GenBank Accession: GCA_000236325.2) Genome Assembly (database) placed the putative ortholog of *DENR* within scaffold KB464954 (KB464954.1) at locus LOC108103491 (XP_017064488.1)— with an E-value of 7e-68 and a percent identity of 51.87%. Furthermore, the putative ortholog is flanked upstream by LOC108103488 (XP_017064485.1) and LOC108103490 (XP_017064487.1), which correspond to *CG4880* and *CG13002* in *D. melanogaster* (E-value: 3e-111 and 1e-101; identity: 52.43% and 67.36%, respectively, as determined by *blastp*; **Figure 1A**, Altschul et al., 1990). The putative ortholog of *DENR* is flanked downstream by LOC108103489 (XP_041675159.1) and LOC108103471 (XP_017064460.1), which correspond to *PolR3I* and *Nprl2* in *D. melanogaster* (E-value: 2e-93 and 0.0; identity: 77.59% and 97.58%, respectively, as determined by *blastp*). The putative ortholog assignment for *DENR* in *D. eugracilis* is supported by the following evidence: the synteny of the genomic neighborhood is completely conserved across both species, and all *BLAST* search results used to determine orthology indicate very high-quality matches.

**Figure 1.**
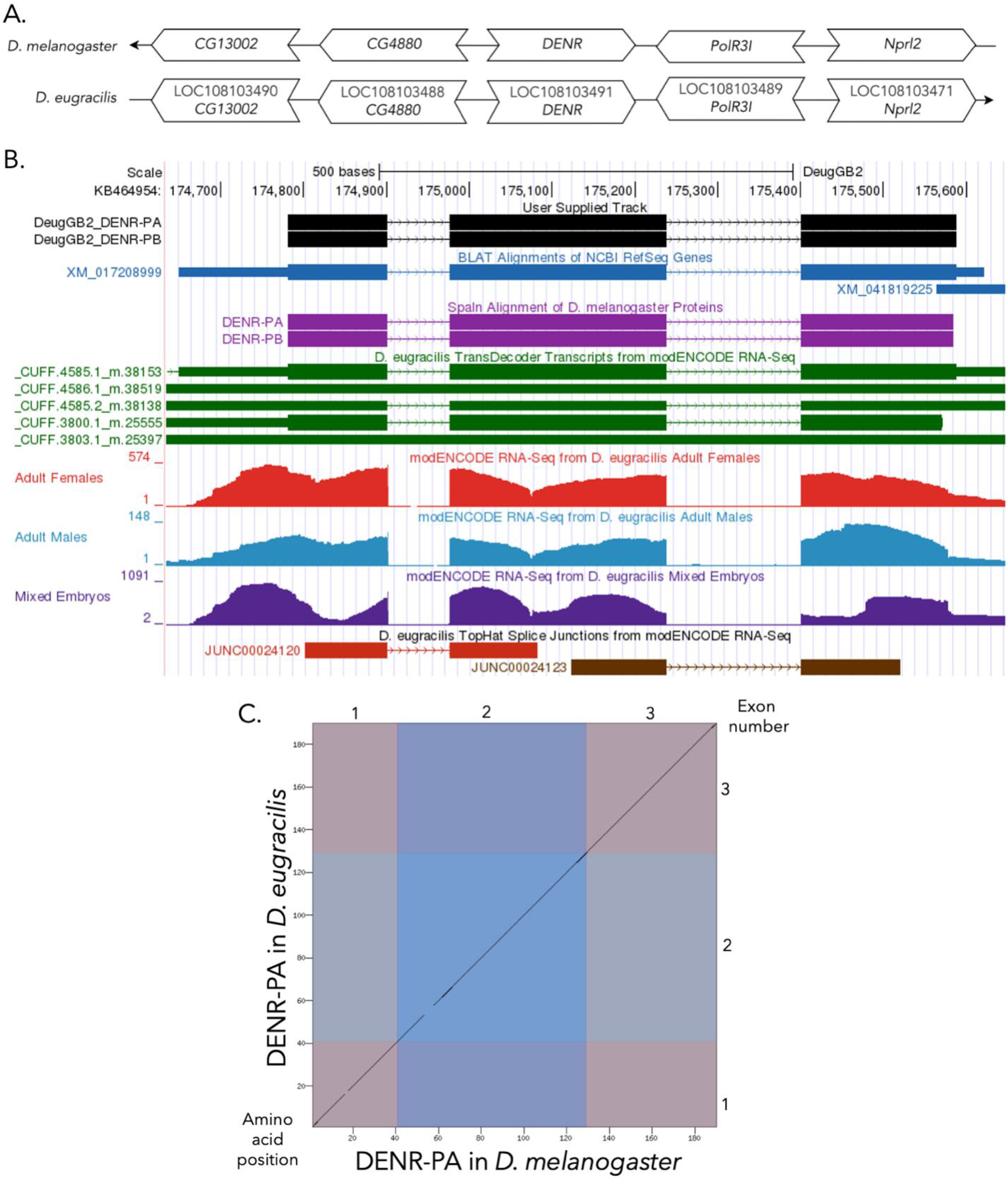
(A) Synteny comparison of the genomic neighborhoods for *DENR* in *Drosophila melanogaster* and *D. eugracilis*. Thin underlying arrows indicate the DNA strand within which the target gene–*DENR*–is located in *D. melanogaster* (top) and *D. eugracilis* (bottom). The thin arrow pointing to the right indicates that *DENR* is on the positive (+) strand in *D. eugracilis*, and the thin arrow pointing to the left indicates that *DENR* is on the negative (-) strand in *D. melanogaster*. The wide gene arrows pointing in the same direction as *DENR* are on the same strand relative to the thin underlying arrows, while wide gene arrows pointing in the opposite direction of *DENR* are on the opposite strand relative to the thin underlying arrows. White gene arrows in *D. eugracilis* indicate orthology to the corresponding gene in *D. melanogaster*. Gene symbols given in the *D. eugracilis* gene arrows indicate the orthologous gene in *D. melanogaster*, while the locus identifiers are specific to *D. eugracilis*. **(B) Gene Model in GEP UCSC Track Data Hub** (Raney et al. 2014). >The coding-regions of *DENR* in *D. eugracilis* are displayed in the User Supplied Track (black); coding exons are depicted by thick rectangles and introns by thin lines with arrows indicating the direction of transcription. Subsequent evidence tracks include BLAT Alignments of NCBI RefSeq Genes (dark blue, alignment of Ref-Seq genes for *D. eugracilis*), Spaln of *D. melanogaster* Proteins (purple, alignment of Ref-Seq proteins from *D. melanogaster*), Transcripts and Coding Regions Predicted by TransDecoder (dark green), RNA-Seq from Adult Females and Adult Males (red and light blue, respectively; alignment of Illumina RNA-Seq reads from *D. eugracilis*), and Splice Junctions Predicted by regtools using *D. eugracilis* RNA-Seq (Chen et al., 2014; PRJNA63469). Splice junctions shown have a minimum read-depth of 769 with 500-999 and >1000 supporting reads in brown and red, respectively. **(C) Dot Plot of DENR in *D. melanogaster* (*x*-axis) vs. the orthologous peptide in *D. eugracilis* (*y*-axis)**. Amino acid number is indicated along the left and bottom; coding-exon number is indicated along the top and right, and exons are also highlighted with alternating colors.

### Protein Model

*DENR* in *D. eugracilis* has two protein-coding isoforms (DENR-PA and DENR-PB; **Figure 1B**). Isoforms DENR-PA and DENR-PB are identical and contain three protein-coding exons. Relative to the ortholog in *D. melanogaster*, the coding-exon number is conserved, as DENR-PA and DENR-PB are also identical with three coding exons in *D. melanogaster*. The sequence of DENR-PA in *D. eugracilis* has 97.88% identity (E-value: 3e-136) with the protein-coding isoform DENR-PA in *D. melanogaster*, as determined by *blastp* (**Figure 1C**). Coordinates of this curated gene model (DENR-PB, DENR-PA) are stored by NCBI at GenBank/BankIt (accession BK064630, BK064631, respectively).

## Methods

Detailed methods including algorithms, database versions, and citations for the complete annotation process can be found in Rele et al. (2023).

## Acknowledgements

We would like to thank Wilson Leung for developing and maintaining the technological infrastructure that was used to create this gene model and Laura K. Reed for overseeing the project.

## Supplemental Files

1. Zip file containing FASTA, PEP, GFF files for the gene model

## Metadata

Bioinformatics, Genomics, *Drosophila*, Genotype Data, New Finding

## References

Altschul, S. F., Gish, W., Miller, W., Myers, E. W., & Lipman, D. J. (1990). Basic local alignment search tool. Journal of Molecular Biology, 215(3), 403–410. 10.1016/S0022-2836(05)80360-2

Bächli, G. (updated 2026, February 4). TaxoDros. TaxoDros. https://www.taxodros.uzh.ch/

Bock, I. R., & Wheeler, M. R. (1972). The Drosophila melanogaster Species Group. In Studies in Genetics VII (pp. 1–102). The University of Texas Publications.

Chen, Z.-X., Sturgill, D., Qu, J., Jiang, H., Park, S., Boley, N., Suzuki, A. M., Fletcher, A. R., Plachetzki, D. C., FitzGerald, P. C., Artieri, C. G., Atallah, J., Barmina, O., Brown, J. B., Blankenburg, K. P., Clough, E., Dasgupta, A., Gubbala, S., Han, Y., … Richards, S. (2014). Comparative validation of the D. melanogaster modENCODE transcriptome annotation. Genome Research, 24(7), 1209–1223. 10.1101/gr.159384.113

Deyo, J. E., Chiao, P. J., & Tainsky, M. A. (1998). Drp, a Novel Protein Expressed at High Cell Density but Not During Growth Arrest. DNA and Cell Biology, 17(5), 437–447. 10.1089/dna.1998.17.437

Duda, O. (1924). Revision der europäischen u. Grönländischen sowie einiger südostasiat. Arten der Gattung Piophila Fallén (Dipteren). Konowia, 3, 97–113.

Fleischer, T. C., Weaver, C. M., McAfee, K. J., Jennings, J. L., & Link, A. J. (2006). Systematic identification and functional screens of uncharacterized proteins associated with eukaryotic ribosomal complexes. Genes & Development, 20(10), 1294–1307. 10.1101/gad.1422006

Gonzalez, J. N., Zweig, A. S., Speir, M. L., Schmelter, D., Rosenbloom, K. R., Raney, B. J., Powell, C. C., Nassar, L. R., Maulding, N. D., Lee, C. M., Lee, B. T., Hinrichs, A. S., Fyfe, A. C., Fernandes, J. D., Diekhans, M., Clawson, H., Casper, J., Benet-Pagès, A., Barber, G. P., … Kent, W. J. (2021). The UCSC Genome Browser database: 2021 update. Nucleic Acids Research, 49(D1), D1046– D1057. 10.1093/nar/gkaa1070

Gramates, L. S., Agapite, J., Attrill, H., Calvi, B. R., Crosby, M. A., Dos Santos, G., Goodman, J. L., Goutte-Gattat, D., Jenkins, V. K., Kaufman, T., Larkin, A., Matthews, B. B., Millburn, G., Strelets, V. B., the FlyBase Consortium, Perrimon, N., Gelbart, S. R., Agapite, J., Broll, K., … Lovato, T. (2022). FlyBase: A guided tour of highlighted features. Genetics, 220(4), iyac035. 10.1093/genetics/iyac035

Grewal, S. S. (2009). Insulin/TOR signaling in growth and homeostasis: A view from the fly world. The International Journal of Biochemistry & Cell Biology, 41(5), 1006–1010. 10.1016/j.biocel.2008.10.010

Hietakangas, V., & Cohen, S. M. (2009). Regulation of Tissue Growth through Nutrient Sensing. Annual Review of Genetics, 43(1), 389–410. 10.1146/annurev-genet-102108-134815

Jenkins VK, Larkin A, Thurmond J. Using FlyBase, a database of Drosophila gnes and genetics. In: Dahmann C, editor. Drosophila: methods and Protocols. New York (NY): Springer; 2022.

Kent, W. J., Sugnet, C. W., Furey, T. S., Roskin, K. M., Pringle, T. H., Zahler, A. M., & Haussler, A. D. (2002). The Human Genome Browser at UCSC. Genome Research, 12(6), 996–1006. 10.1101/gr.229102

Larkin, A., Marygold, S. J., Antonazzo, G., Attrill, H., dos Santos, G., Garapati, P. V., Goodman, J. L., Gramates, L. S., Millburn, G., Strelets, V. B., Tabone, C. J., Thurmond, J., FlyBase Consortium, Perrimon, N., Gelbart, S. R., Agapite, J., Broll, K., Crosby, M., Dos Santos, G., … Lovato, T. (2021). FlyBase: Updates to the Drosophila melanogaster knowledge base. Nucleic Acids Research, 49(D1), D899–D907. 10.1093/nar/gkaa1026

Laskowski, L. F., Burton, I., Stanek, T. J., Findlay, G. D., Tanner, S., Vincent, J. A., Chak, S. T., Ellison, C. E., & Rele, C. P. (2024). Gene model for the ortholog of DENR in Drosophila yakuba. 10.17912/micropub.biology.001017

Morgan, A., Kiser, C. A., Bronson, I., Lin, H., Guillette, N., McMahon, R., Kennell, J. A., Long, L. J., Reed, L. K., & Rele, C. P. (2002). Drosophila eugracilis – Akt. microPublication Biology. 10.17912/micropub.biology.000544

Mudge, J. M., & Harrow, J. (2016). The state of play in higher eukaryote gene annotation. Nature Reviews Genetics, 17(12), 758–772. 10.1038/nrg.2016.119

Myers A., Hoffmann A., Natysin M., Arsham A.M, Stamm J., Thompson J.S., Rele C.P. 2024. Gene model for the ortholog Myc in Drosophila ananassae, microPublication Biology, submitted.

Pélandakis, M., & Solignac, M. (1993). Molecular phylogeny of Drosophila based on ribosomal RNA sequences. Journal of Molecular Evolution, 37(5), 525–543. 10.1007/BF00160433

Raney, B. J., Dreszer, T. R., Barber, G. P., Clawson, H., Fujita, P. A., Wang, T., Nguyen, N., Paten, B., Zweig, A. S., Karolchik, D., & Kent, W. J. (2014). Track data hubs enable visualization of user-defined genome-wide annotations on the UCSC Genome Browser. Bioinformatics, 30(7), 1003–1005. 10.1093/bioinformatics/btt637

Rele, C. P., Sandlin, K. M., Leung, W., & Reed, L. K. (2023). Manual annotation of Drosophila genes: A Genomics Education Partnership protocol. F1000Research, 11, 1579. 10.12688/f1000research.126839.3

Schleich, S., Strassburger, K., Janiesch, P. C., Koledachkina, T., Miller, K. K., Haneke, K., Cheng, Y.-S., Küchler, K., Stoecklin, G., Duncan, K. E., & Teleman, A. A. (2014). DENR–MCT-1 promotes translation re-initiation downstream of uORFs to control tissue growth. Nature, 512(7513), 208–212. 10.1038/nature13401

Skabkin, M. A., Skabkina, O. V., Dhote, V., Komar, A. A., Hellen, C. U. T., & Pestova, T. V. (2010). Activities of Ligatin and MCT-1/DENR in eukaryotic translation initiation and ribosomal recycling. Genes & Development, 24(16), 1787–1801. 10.1101/gad.1957510

Tello-Ruiz, M. K., Marco, C. F., Hsu, F.-M., Khangura, R. S., Qiao, P., Sapkota, S., Stitzer, M. C., Wasikowski, R., Wu, H., Zhan, J., Chougule, K., Barone, L. C., Ghiban, C., Muna, D., Olson, A. C., Wang, L., Ware, D., & Micklos, D. A. (2019). Double triage to identify poorly annotated genes in maize: The missing link in community curation. PLOS ONE, 14(10), e0224086. 10.1371/journal.pone.0224086

